# Hot-starting software containers for bioinformatics analyses

**DOI:** 10.1101/204495

**Authors:** Pai Zhang, Ling-Hong Hung, Wes Lloyd, Ka Yee Yeung

**Author notes:** Correspondence should be addressed to K.Y.Y.

## Abstract

Using software containers has become standard practice to reproducibly deploy and execute biomedical workflows on the cloud. We demonstrate that hot-starting, from containers that have been frozen after the application has already begun execution, reduces the costs of cloud computing by avoiding repetitive initialization steps. The method is widely applicable and can provide substantial savings both for small jobs and for large-scale deployments using automated schedulers.

---

With the availability of high-throughput next generation sequencing technologies and the subsequent explosion of big biomedical data, the processing of biomedical big data has become a major challenge. Cloud computing plays an important role in addressing this challenge by offering massive scalable computing and storage, data sharing and on-demand access to resources and applications ^1,2^. The National Institutes of Health is launching a Data Commons Pilot Phase to provide access and storage of biomedical data and bioinformatics tools on the cloud (https://commonfund.nih.gov/bd2k). Additionally, software containers have become increasingly popular for deploying bioinformatics workflows on the cloud. Docker (https://www.docker.com/), an open source project, has become the *de facto* standard for container software. Docker packages executables with all the necessary software dependencies ensuring that the same software environment is replicated regardless of the host hardware and operating system. Thus, containerization enhances the reproducibility of bioinformatics workflows. In the context of cloud computing, the utility of containers comes from the ease in which a virtual cloud cluster can be rapidly provisioned with the all the necessary dependencies for a complicated workflow by simply downloading a set of containers, each of which take a few seconds to spin up. Recently, Vivian et al. processed over 20,000 RNA sequencing (RNA-seq) samples from the Cancer Genome Atlas (TCGA) using Docker containers on the cloud ^3^. Tatlow *et al.* used software containers to study the performance and cost profiles of different cloud-based configurations in processing RNA-seq data from public cancer compendia ^4^.

When containers are deployed, applications are launched *de novo* each time the container is spun up. This means that any initial preparatory steps are repeated each time the container is used. For applications such as the alignment of reads, these initial steps can be quite substantive as a entire reference genome is read in and indices are generated. In an automated large-scale deployment, these steps are replicated many times. It would be far more efficient if one could “*checkpoint*” and save containers in states where the application has already completed the initialization steps so as to avoid unncessary repetitions. One could then “*hot-start*” workflows from these checkpoints. This is analogous to hot-start PCR where all the necessary reagents are pre-mixed awaiting only the addition of the template.

Our key idea is to save and restore memory states in software containers using the Checkpoint Restore in Userspace (CRIU) tool. CRIU freezes a running container and saves the checkpoint as a collection of files on disk (https://criu.org/Main_Page). These files can subsequently be used to restore and resume the application from that checkpoint. CRIU was originally developed for Linux, but has recently become available for Docker (https://criu.org/Docker). While it is possible to stop Docker containers with native docker commands, this process does not preserve the memory state. Although re-starting from a ready-to-go state is an intuitive application of checkpointing, we have been unable to find any previous description of using checkpointing as a general method for improving the efficiency of container deployments.

We demonstrate that hot-starting from a saved container checkpoint can significantly reduce the execution time using the STAR aligner ^5, 6^ for RNA-seq data analyses. We choose STAR as a proof-of-concept example because it has an option to save an intermediate state. However, our idea of using checkpoints has broad applications in optimizing performance using software containers on the cloud *when deploying any bioinformatics task where a pause could be inserted to capture a re-usable state*.

The STAR aligner ^5, 6^ consists of two steps. In the first step, genome indices using the reference genome as input are generated. In the second step, read sequences from a specific experiment sample are mapped to the reference genome assuming that the genome indices have already been generated. In particular, STAR has the option of keeping the indices in memory after they have been generated to avoid repeating the first step when multiple files are to be aligned to the same reference genome. We used the CRIU tool to create checkpoints after the first step of generating genome indices. Instead of launching a new container and starting STAR from scratch, we restore the container state using CRIU and resume running STAR after it has loaded the indices. Figure 1 shows an overview of our approach with and without using checkpoints.

**Figure 1.**
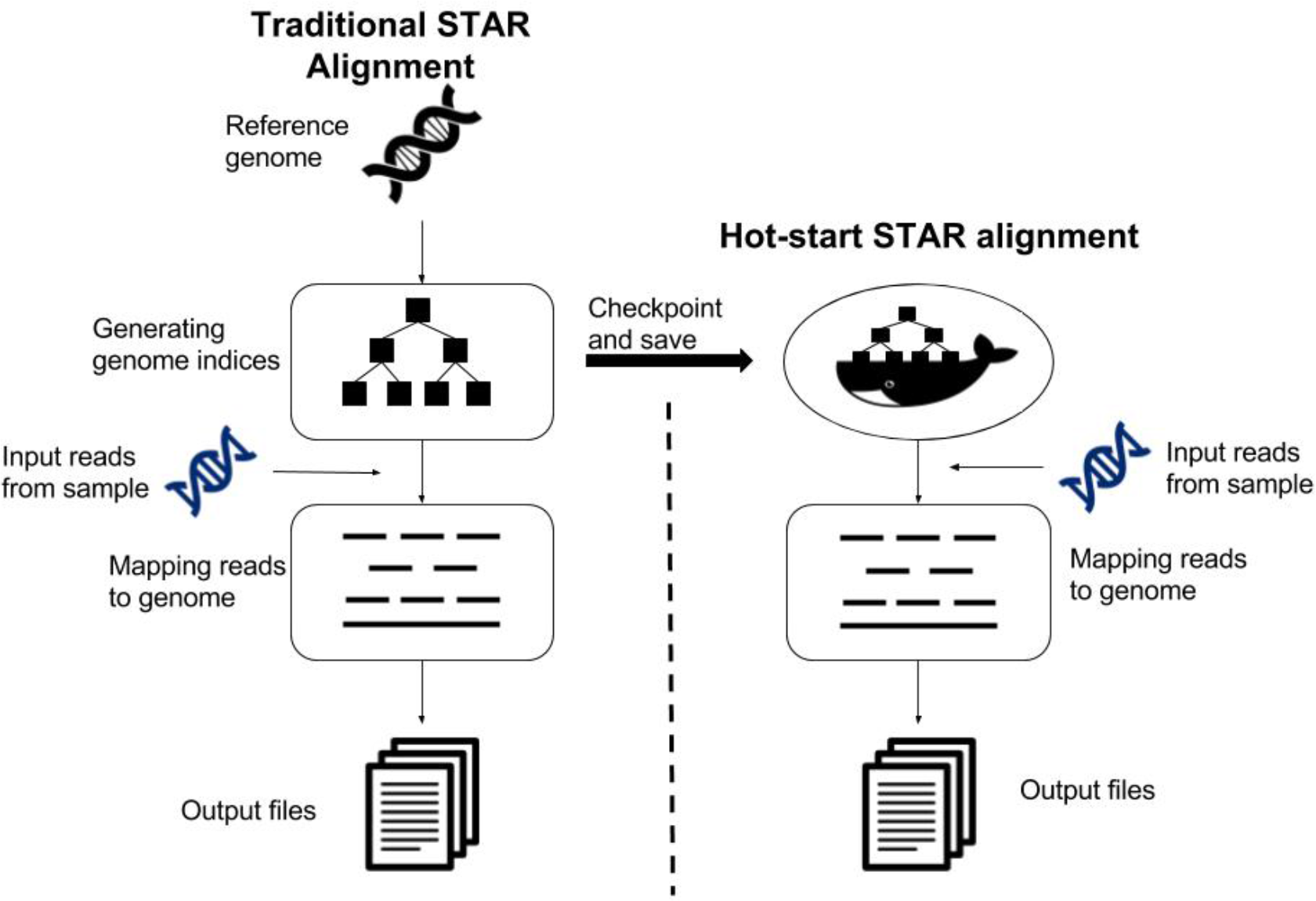
An overview of our approach with and without checkpoints. The left panel shows the two steps of the STAR aligner ^5, 6^. Note that the generation of the genome indices in the first step only requires the reference genome as input, thus the data from the experimental sample is only used in the second (mapping) step of STAR. The right panel shows our approach using the Checkpoint Restore in Userspace (CRIU) tool that freezes a running container and saves the checkpoint as a collection of files on disk after the genome indices are generated using the reference genome. Our “hot-start” containers use these saved files to restore the application and map the reads from the experimental sample data to the reference.

To test the checkpointing methodology, we used RNA-seq data generated by Himes *et al.* which measure the gene expression changes in human airway smooth muscle cells in response to asthma medications ^7^. We compared the time required to align the sequences with a normal container where STAR starts from scratch, and the time required when hot-starting from a container checkpoint where STAR has already generated indices. We performed empirical studies on cloud instances including the Amazon Web Services (AWS) and Microsoft Azure, using both local and networked disks. On AWS, we compared the performance with data on the local host and Amazon Elastic Block Store (EBS). On Microsoft Azure, we compared the performance with data on local host and Azure File Storage. Please refer to Methods for details of our experimental setup. Our empirical results are shown in Figure 2.

**Figure 2.**
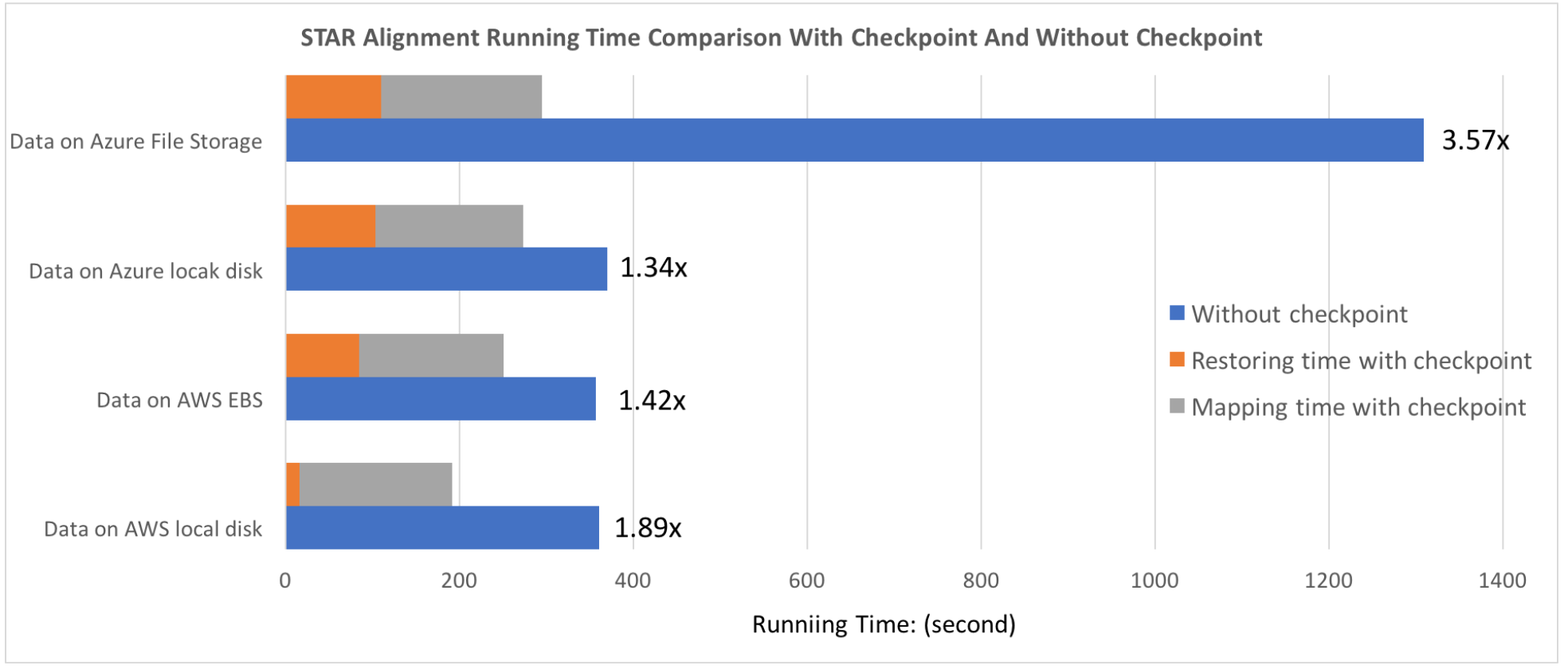
STAR alignment running time comparison with checkpoint and without checkpoint. We performed our empirical experiments on two cloud platforms: Amazon Web Services (AWS) and Microsoft Azure. Both the Azure File Storage and the Amazon Elastic Block Store (EBS) represent network disks. We observe that our “hot-start” containers (orange and grey bars) provide major reduction in execution time, especially on local disks.

Figure 2 shows that the STAR aligner with checkpoint reduces the execution time compared to STAR without checkpoint. On AWS, we observed 1.89x speedup with data stored on the local disk and 1.42x speedup with data on network disk (Amazon EBS). On Microsoft Azure, we achieved a 1.34x speedup with data stored on the local disk, and 3.57x speedup with data on Azure File Storage. With respect to execution time, we show that hot-starting from checkpoint containers save 2 minutes on fast local disks and Amazon EBS disks. The saving is almost 20 minutes when using Azure network storage where the disk caching scheme appears to be much less favorable to STAR’s index generation process.

In this article, we have presented a novel idea for optimising cloud deployments using checkpointing to save containers where the applications are already started. Using CRIU for Docker, we can save the container with a preloaded genome for STAR alignment and restore the container from these checkpoint files to any host. We have achieved successful migration of checkpointed containers to different virtual machine instances running on the Amazon and Azure cloud platforms while realizing up to a 3.57x speedup using our approach saving up to 20 minutes for a single STAR alignment workflow on Azure with network disks. For STAR alignment, it is possible to use a checkpointed container to align multiple sequences at once by retaining the genomic indices in memory. Our approach yields a significant benefit with hot-starting when as few as one or two files are aligned. Additionally, multiple STAR alignment tasks can be computed in parallel using the same genome indices hosted by different processes. For automated schedulers such as Docker Compose (https://docs.docker.com/compose/), “hot-starting” reduces execution time every single time the STAR container is launched. While it is possible to design a workflow to perform all the alignments in a single container first, load-balancing optimizations would be better utilized by allowing the scheduler to distribute the computation over the cluster as shorter jobs.

Our hot-start strategy only requires that there is a convenient place for a pause, checkpoint and re-start. In the case of STAR, this is provided by a flag that allows the container to keep genomic indices in shell memory between invocations of STAR. For other workflows, one could add a flag to pause the computation where the checkpoint is to be created, and a flag to resume the computation afterwards. With these straightforward modifications, any application could take advantage of checkpointing to avoid repetitive initialization. This is a novel and unexplored approach to optimising containerized workflows while reducing the costs of cloud computing.

## ACKNOWLEDGEMENTS

L.H.H. and K.Y.Y. are supported by NIH grant U54HL127624 and the AMEDD Advanced Medical Technology Initiative. We would like to acknowledge support from the AWS Cloud Credits for Research (to Lloyd and Yeung) and the Microsoft Azure for Research programs (to Hung and Lloyd) for providing cloud computing resources. We would like to acknowledge the Student High Performance Computing Club and the eScience Institute at University of Washington for both technical assistance and computing resources to Pai Zhang.

## AUTHOR CONTRIBUTIONS

P.Z. implemented the Docker containers and conducted the empirical experiments. P.Z., L.H.H. and K.Y.Y. drafted the manuscript. K.Y.Y. and L.H.H. designed the case study. W.L. provided cloud computing expertise. K.Y.Y. coordinated the empirical study. All authors edited the manuscript.

## METHODS

**CRIU.** CRIU (Checkpoint/Restore In Userspace) is a Linux software tool that freezes a running application and saves it as collection of files to disk (https://criu.org). The application can later be restored on the same or on a different host. Docker currently integrates CRIU as an experimental checkpoint sub-command that saves the state of processes to a collection of files on disk. The checkpointing command has been used to migrate containers from the source host to target host when the resources of the source are limited ^8^, for fault tolerance purposes ^9^, and can provide highly available and scalable of micro-services^10^.

### Cloud configurations tested

In our experiments, we deployed our containers on instances from two cloud platforms:
Microsoft Azure and Amazon Web Services (AWS). Ubuntu 16.04 was the host operating system in all our tests. Testing was conducted using a standard DS13 v2 instance with 8 virtual CPUs and 56 Gb memory on a Azure and a m4.4xlarge instance with 16 virtual CPUs and 64 Gb memory on AWS. As disk I/O is an important factor in the efficiency of CRIU restoration and the generation of indices without CRIU, instances were tested using both network based disks (EBS for AWS and Microsoft Azure File Storage for Azure) and locally attached disks.

### Creating hot-start containers

We installed CRIU on the host Ubuntu system. Docker Community Edition (Docker CE), which includes the experimental checkpointing tool, was then installed. The STAR binary was compiled from source (https://github.com/alexdobin/STAR) using Ubuntu 16.04 and g++ and then copied into a fresh Ubuntu 16.04 container to get rid of all the intermediate build files. The build code and Dockerfiles are available from https://github.com/BioDepot/ubuntu-star. To create the checkpoint, STAR was launched with the *genomeLoad* flag set to *LoadAndKeep.* This keeps the indices in shared memory after STAR exits. To trap the container in this state, we launched STAR using a parent shell script that did not exit, and checkpointed the container after STAR exited. This results in the generation of checkpoint files that store the state of the hot-start container. Due to different kernel versions being used, we created separate hot-start containers for AWS and Azure.

### Comparing hot-start containers and standard cold-start containers

The paired-end fastq files were 9 Gb in size comprising 22935521 reads. Times were recorded for the generation of aligned BAM files using STAR in the standard container and using STAR with the hot-start container. Times include the time required to restore the hot-start container from the checkpointed files.

**Software availability.** The scripts we used to deploy the containers in our empirical study are publicly available on GitHub and the Docker images are available on DockerHub. GitHub URL: https://github.com/paizhang/Hotstarting-For-STAR-Alignment DockerHub URL: https://hub.docker.com/rZbiodepot/star-for-criu/

**Data availability.** The fastq files used in our tests were generated by Himes *et al.* and is publicly available from GEO with accession number GSE52778 (https://www.ncbi.nlm.nih.gov/geo/query/acc.cgi?acc=GSE52778).

